# Self-distillation and antisymmetric constraints enable accurate prediction of protein stability changes from single mutations

**DOI:** 10.1101/2025.05.18.654422

**Authors:** Wenkang Wang, Min Li

## Abstract

Accurately predicting protein stability changes upon residue-level mutations is essential for protein engineering and design. However, previous computational methods are typically designed for specific tasks and struggle to generalize across different stability systems, preventing the accurate guidance of protein mutagenesis. Sequence-based methods are easier to apply to large-scale proteins since they do not rely on high-quality structures. However, existing sequence-based approaches struggle to capture structural changes, resulting in lower performance compared to structure-based methods. In this study, we propose DeepTherm, a sequence-based deep learning solution to model changes in folding free energy (ΔΔ*G*) and melting temperature (Δ*T*_*m*_) resulting from single-point mutations. DeepTherm transfers a protein large language model as a core component and incorporates a cross-attention mechanism to capture the contact-map changes around mutated positions. To address data imbalance and the antisymmetric nature of mutation effects, DeepTherm employs a self-distillation inference strategy under the supervision of an antisymmetric constraint. Benchmarking across multiple datasets shows that DeepTherm achieves state-of-the-art accuracy in both ΔΔ*G* and Δ*T*_*m*_ predictions. Furthermore, practical evaluations confirm DeepTherm’s capability in accurately ranking protein stability on large-scale datasets and effectively identifying critical structural contacts impacting stability. Detailed analysis on cDNA display proteolysis data further highlights the critical importance of self-distillation and antisymmetric constraints in enhancing predictive performance. The source code can be found in https://github.com/CSUBioGroup/DPStab.

## 1 Introduction

The thermal stability of a protein is essential for maintaining its biological functions in living organisms [1, 2]. Protein stability is closely linked to its amino acid sequence and three-dimensional structure, both of which can be significantly altered by single residue mutations, thereby disrupting the overall stability and potentially impairing functions [3, 4]. Consequently, introducing specific mutations to enhance protein stability has become a common strategy, particularly in industrial and therapeutic protein engineering [5, 6]. However, brute-force mutagenesis approaches are both time-consuming and costly. Therefore, developing computational methods for accurately predicting protein stability changes upon residue mutations can reveal crucial insights into the relationships between protein sequence, structure, stability, and function, offering valuable guidance for rational protein design [7].

In the context of protein stability change prediction, stability is typically quantified by the Gibbs free energy (Δ*G*) and melting temperature (*T*_*m*_). Specifically, Δ*G* represents the free energy difference between the folded and unfolded states (*G*_*unfolded*_ − *G*_*folded*_) and *T*_*m*_ denotes the midpoint of thermal denaturation. Correspondingly, stability changes upon mutation are commonly expressed as changes in Gibbs free energy (ΔΔ*G*) and melting temperature (Δ*T*_*m*_), which are both antisymmetric: ΔΔ*G*_*x→y*_ = −ΔΔ*G*_*y→x*_ and Δ*T*_*m,x→y*_ = −Δ*T*_*m,y→x*_. In this study, higher ΔΔ*G* and Δ*T*_*m*_ indicate stabilizing mutations.

To accurately predict ΔΔ*G* and Δ*T*_*m*_, numerous predictors centered on the geometric potential energy of protein structures have been proposed, such as FoldX [8], EvoEF [9, 10], STRUM [11], and Korpm [12]. These computational methods rely on protein structures, rendering them limited when high-quality experimental or predicted structures are not available. Additionally, machine learning methods, like DDGun [13, 14] and Mutate Everything [15], have also been proposed for ΔΔ*G* prediction from sequences. Notably, a few deep learning approaches [16, 17] integrate protein sequence and structure information to predict ΔΔ*G* and Δ*T*_*m*_, which achieve an improvement compared to other methods and provide new insights for incorporating sequence and structure information through deep learning. However, these methods require the native or predicted protein structures, which prevents them from performing end-to-end prediction directly from sequence. Meanwhile, the imbalance of protein stability data presents a significant challenge for existing methods, where the number of stabilizing mutations is considerably less than that of destabilizing mutations. This phenomenon can easily impact these approaches, leading to results with distinct preferences [18].

Protein large language models (pLLMs) [19–21], owing to their self-supervised training on massive unlabeled protein data, exhibit remarkable generalization capabilities and have achieved robust performance across numerous downstream tasks [2, 19]. Recently, some efforts have attempted to use pLLMs to predict protein stability changes [22, 23]. For instance, ThermoMPNN [23] extracts the learnable features from a structure-based pLLM, ProteinMPNN [24], and leverages transfer learning from the mega-scale data used in ProteinMPNN to the protein stability data. In contrast, PROSTATA [22] adopts a sequence-based pLLM, ESM2 [20], as the backbone and uses different regression heads to predict protein stability changes. However, most existing pLLMs and structure modeling methods treat each protein in isolation during pre-training, which limits their ability to distinguish the single mutation effects. Although the overall protein backbone may remain globally unchanged upon single mutations, the subtle yet crucial alterations in local residue contacts around mutation sites are often critical determinants of stability. These local impacts are challenging to capture for models that primarily rely on global protein information.

Consequently, it is essential to capture the subtle contact changes around the mutation sites, even when the overall protein backbone topology remains largely unaltered. Additionally, distillation on existing LLMs shows great potential to enhance the performance of LLMs for specific downstream tasks [25] and has made much progress in various areas [26], such as DeepSeek-R1 [27], providing new perspectives for LLM transfer. Here, we propose DeepTherm, a protein sequence-based deep learning method for accurate prediction of protein stability changes. A key strength of DeepTherm lies in its ability to extract residue-level features, infer contact patterns, and integrate local contextual information around mutation sites within the architecture of a protein LLM. To further enhance predictive accuracy, we introduce a self-distillation inference strategy and an antisymmetric constraint tailored to the nature of mutation effects. Comprehensive benchmarks and data analyses demonstrate that DeepTherm achieves state-of-the-art accuracy for both ΔΔ*G* and Δ*T*_*m*_ prediction.

## 2 Methods

The overall DeepTherm pipeline is illustrated in Fig. 1a. Starting from a pair of proteins (wild-type and mutated protein sequences), DeepTherm first leverages a set of transformer layers to extract comprehensive evolutionary and conformational information from both the wild-type and mutated protein sequences (Fig. 1b). These layers process each sequence separately, generating their respective residue representations and inferring their contact maps. DeepTherm then introduces a neighboring encoder to capture the local structural changes around the mutation site (Fig. 1c). It cross-incorporates information between the residue representations and inferred contact information, assigns weighted attention to neighboring residues, and adaptively integrates neighboring information. Finally, DeepTherm predicts thermal stability changes through a feature transition module (Fig. 1d), which compares the wild-type and mutant protein representations and incorporates external factors to predict ΔΔ*G* or Δ*T*_*m*_.

**Fig. 1.**
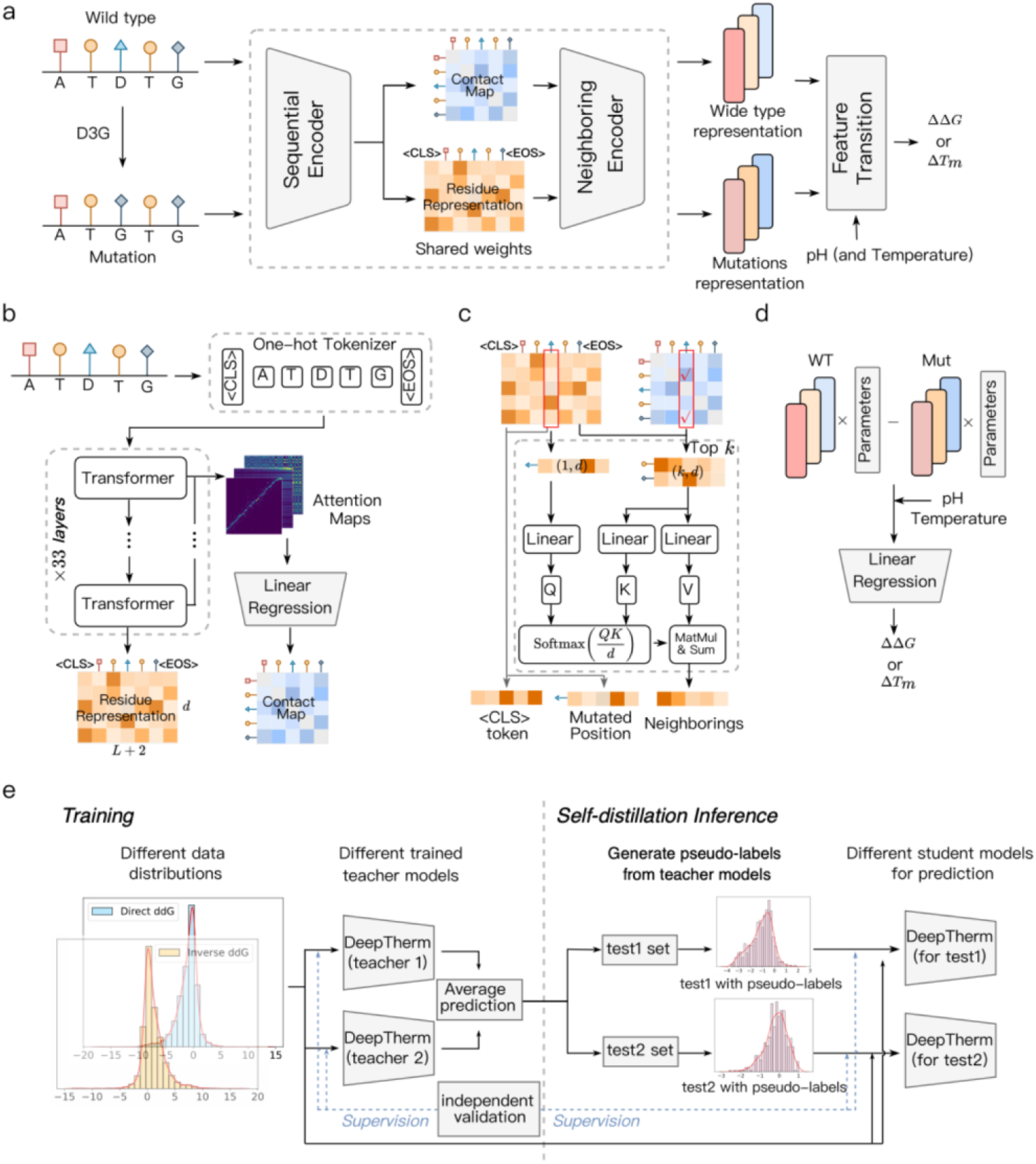
Model architecture of DeepTherm. **a**. The overview of DeepTherm. It consists of three modules. The wild-type protein sequence and mutated protein sequence are fed into the first two modules, respectively, to generate the features of the whole protein, mutated position, and its neighboring information. Subsequently, the features of wild-type and mutated proteins are combined in a feature transition module, with environmental parameters (pH and temperature), to predict the difference of stability (Δ*G* or Δ*T*_*m*_) between two proteins. **b**. The architecture of the sequential encoder module. **c**. The architecture of the neighboring encoder module. **d**. The details of the feature transition module. **e**. The details of self-distillation for different test datasets.

Additionally, at the inference stage, we propose a self-distillation strategy to enrich the training data (Fig. 1e) and incorporate a constraint to align DeepTherm with the antisymmetric nature of mutations. Specifically, multiple DeepTherm models are first trained separately on training data with different distributions. These trained models act as teachers to produce pseudo-labels for test data, which will be fed back to generate better student models with the original training set and independent validation set. Meanwhile, a contrastive loss is applied to make DeepTherm produce opposite outputs for direct and inverse mutations. Further details of each module can be found in the “Methods” section.

### 2.1 Architecture of sequential encoder

To learn evolutionary and conformational information of protein sequence, DeepTherm adopts the same sequential encoder as ESM2 and utilizes the pre-trained weights of ESM2 as its initial parameters (Fig. 1b), including one token embedding layer, 33 stacked transformer layers, and one linear regression layer for contact map prediction. Specifically, the input is a wild type or mutated protein sequence *p* = {*r*_1_, *r*_2_, …, *r*_*L*_} with *L* residues, where *r*_*i*_ ∈ {*A, C*, …, *Y*} are the different residue types. It is first fed into a token embedding layer:

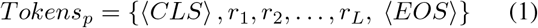

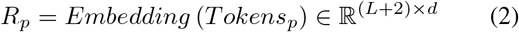

where *Tokens*_*p*_ is the tokens that represent the protein sequence *p*, ⟨*CLS*⟩ and ⟨*EOS*⟩ are two additional signs indicating the beginning and end of protein sequences, and *d* is the representation dimension. Then, a trainable embedding matrix is used to generate corresponding representations for each token, denoted as *R*_*p*_, which is fed into later transformer layers:

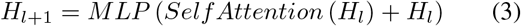

where *H*_*l*_ ∈ ℝ ^(*L*+2)*×d*^ is the input of (*l* + 1) − *th* transformer layer (*l* = 0, …, *n* − 1), and *H*_0_ = *R*_*p*_ initially. *Self Attention*(•) is traditional attention mechanism that generates (*Q, K, V*) items, calculates attention scores between different tokens and updates their representations. *MLP* (•) is composed of several linear layers and corresponding activate functions. Through transformer layers, different attention maps can be obtained from different heads, which are concatenated to predict contact probabilities between residues:

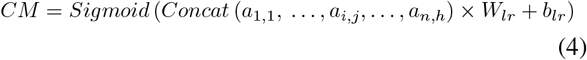

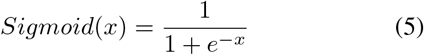

where *α*_*i,j*_ is the attention map from *j* − *th* attention heads in *i* − *th* transformer layer, (*W*_*lr*_ ∈ ℝ ^(*n* × *h,1)*^, *b*_*lr*_ ∈ ℝ^1^) is a linear layer, and (*n, h*) represent the number of transformer layers and attention heads of each layer, respectively.

### 2.2 Architecture of neighboring encoder

Through sequential encoder, DeepTherm generates the residue representations *H*_*n*_ and corresponding contact map *CM* from the target protein sequence *p*. Based on these features, DeepTherm introduces a cross-attention module to extract the neighboring information of mutated site *m* (1 ⩽ *m* ⩽ *L*). As shown in Fig. 1c, in the contact map *CM*, the top-k neighboring residues with the highest contact probabilities with the mutation site *m* are selected:

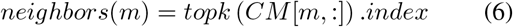

Subsequently, the representations of these *k* neighbors (denoted as *H*_*n*_ [*neighbors*(*m*), :] ∈ ℝ^*k×d*^), along with the mutation site (*H*_*n*_[*m*, :] ∈ ℝ^1*×d*^), are then extracted and used as inputs for the cross-attention module to integrate the neighboring information:

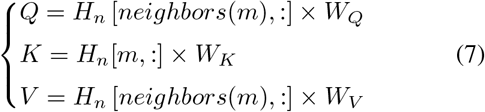

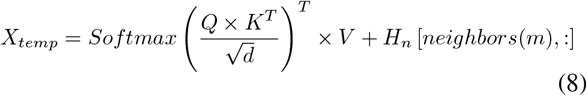

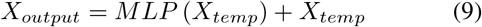

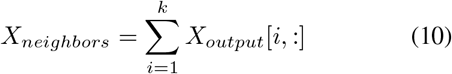

where (*Q* ∈ ℝ^*k×d*^, *V* ∈ ℝ^*k×d*^) is linearly transferred from the *k* neighbors and *V* ∈ ℝ^1*×d*^ is from the mutated site. Through this cross-attention mechanism, DeepTherm can learn the importance of neighbors and integrate their representations adaptively, denoted as *X*_*neighbors*_ ∈ ℝ^1*×d*^. Additionally, residual operation and *LayerNorm*(•) are also employed here to prevent the loss of features.

### 2.3 Architecture of feature transition module

Finally, by two previous encoders, DeepTherm generates the features of the target protein, mutated site, and corresponding neighbors, denoted as *H*_*n*_[0, :] ∈ ℝ^1*×d*^ (⟨*CLS*⟩ token), *H*_*n*_[*m*, :] ∈ ℝ^1*×d*^, and *X*_*neighbors*_ ∈ ℝ^1*×d*^, respectively. Hence, the features of wild-type protein 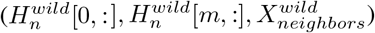 and mutated protein 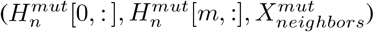 can be obtained separately. Based on these paired features, a simple feature transition module, as illustrated in Fig. 1d, is applied to learn the difference between the two proteins:

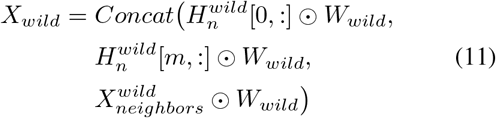

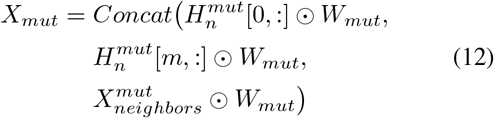

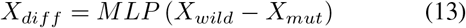

where *W*_*wild*_ ∈ ℝ^1*×d*^ and *W*_*mut*_ ∈ ℝ^1*×d*^ are two learnable parameters. After obtaining the difference between wild-type and mutated proteins, environmental parameters are combined to predict Δ*G* changes (ΔΔ*G*) upon single-point mutation:

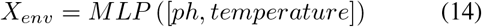

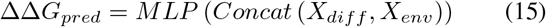

### 2.4 Self-distillation strategy and model training

As shown in Fig. 1e, the whole process of DeepTherm can be divided into two stages, including training and inference. Specifically, the first stage generates two different trained DeepTherm models, based on the direct mutations and symmetrical mutations. To effectively learn the naturally imbalanced distribution dominated by destabilizing mutations, the first DeepTherm model is trained on direct training data and optimized using mean squared error (MSE) loss function:

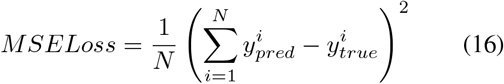

where *N* is the number of training samples and 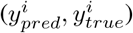 are the predicted and true ΔΔ*G* values, respectively. Then, we train another DeepTherm model on symmetrical data with MSE loss and an additional antisymmetric constraint:

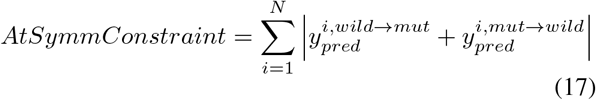

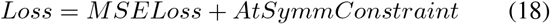

Where 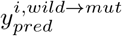 is the predicted result of *i* − *th* training sample from wild-type protein to mutated protein, and 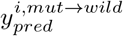 corresponds to the predicted result for the inverse mutation. This loss function ensures that the model’s predictions are consistent with the constraint as much as possible. Based on these two teacher models, at the inference stage, we first average their predicted results for test data as pseudo-labels. After incorporating training data and test data with pseudo-labels, a single dependent DeepTherm is trained to produce the final predictions. In the entire training and inference process, the hidden dimension size is set to 1280, AdamW optimizer is used to optimize models with dynamic decaying learning rates, and the weights of transformer layers from ESM2 are fixed in the first two epochs. DeepTherm will stop training if the loss value of validation data is stable or trending upwards.

All experiments of DeepTherm are carried out using one NVIDIA Tesla H100 GPU card with 96 GB of memory. The training process takes around 15 hours with a batch size of 4.

## 3 Experiments and results

### 3.1 Datasets

The ΔΔ*G* and Δ*T*_*m*_ training dataset used in this article is derived from the previous study [16], named S8754 and S4346, respectively, which are collected from the ProThermDB [28] and ThermoMutDB [29] databases. S8754 consists of 8754 single-point mutations across 301 proteins, along with environmental factors (pH and temperature) and ΔΔ*G* values. S4346 includes 358 proteins with 4346 single residue mutations, along with pH factors and Δ*T*_*m*_ values. We randomly select 10% of the training data to serve as a validation set to assess whether the model training is sufficient. Additionally, we use an independent dataset to test the performance of models in ΔΔ*G* prediction, named S461. This dataset maintains *<*25% sequence identity with the proteins in S8754 and each mutation has been manually corrected. Finally, there are 461 single mutations of 48 proteins used in our test dataset. For Δ*T*_*m*_ prediction, we use S571 as test data. It includes 39 proteins with 571 single mutations, which are all controlled by 25% sequence identities to the S4346.

Due to the antisymmetric nature of mutations, where ΔΔ*G*_*wild→mut*_ = −ΔΔ*G*_*mut→wild*_, it is necessary to measure the prediction results of models in antisymmetric consistency. Consequently, we generate corresponding inverse mutation data (‘mutation’→’wild type’) from the direct mutation data (‘wild type’→’mutation’), resulting in a maximum of 17235 samples available for training and 922 samples for test. Similarly, there are 8692 samples for Δ*T*_*m*_ training and 1142 samples for test.

Moreover, we also collect an inferred ΔΔ*G* dataset generated by cDNA display proteolysis [30], consisting of 389068 single residue mutations across 412 proteins. Based on the length of the wild-type protein sequences, we randomly select three proteins within each length interval of 7 residues and randomly selected 150 mutations per protein to construct the test set. Totally, there are 6 generated test sets, each containing approximately 450 mutations.

### 3.2 Evaluation metrics

In this study, seven metrics are used to evaluate the performance of models in regression and classification, including Spearman correlation coefficient (SCC), Pearson’s correlation coefficient (PCC), root mean square error (RMSE), and mean absolute error (MAE), accuracy (ACC), Matthews correlation coefficient (MCC) and Cohen’s kappa score. Specifically, given predicted results *Y*_*pred*_ and true labels *Y*_*true*_, SCC is defined as:

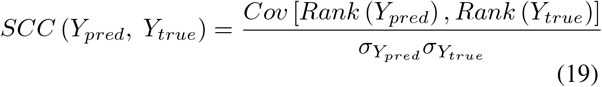

where *Rank*(•) is the rank positions of elements in the sorted order of the original list. *Cov* denotes the covariance between two lists, and *σ* is the standard deviation. PCC is similar to SCC but directly calculates the correlation between the original lists:

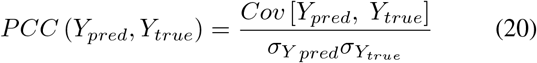

where larger SCC and PCC indicate better predictions. RMSE and MAE reflect the error distances between predictions and labels:

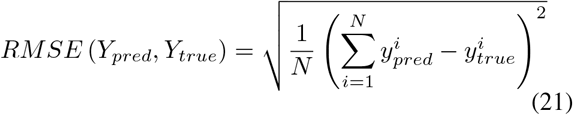

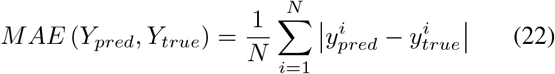

Both smaller RMSE and MAE represent better performance. Additional three metrics, ACC, MCC, and Cohen’s kappa score (*κ*) evaluate the multi-class classification abilities of models:

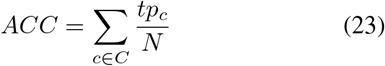

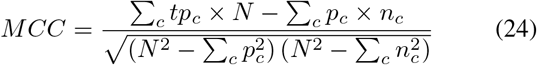

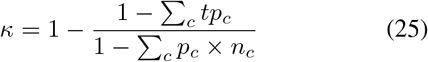

where *tp*_*c*_ denotes the number of true positives for category *c*, and *N* represents the number of samples. *p*_*c*_ and *n*_*c*_ represent the number of samples predicted as class *c* and the number of samples belonging to class *c*, respectively. Both higher ACC, MCC, and *κ* values show better performance in multi-class classification.

### 3.3 DeepTherm outperforms existing methods overall in ΔΔG and ΔT_m_ prediction

To fairly evaluate the overall performance of DeepTherm in ΔΔ*G* prediction, we utilize a commonly used manually curated independent test dataset, S461 [12], containing proteins sharing a low sequence identity with those in the training set. S461 was also used as a benchmark set in previous studies [16, 17]. We further benchmark DeepTherm against seven structure-based (FoldX [8], DDGun-3D [13, 14], GeoDDG-3D [16], GeoDDG-AF2 [16], ThermoMPNN [23], mCSM [31], ThermoNet [32]) and three sequence-based (DDGun [13, 14], Mutate Everything [15], PROSTATA [22]) approaches. We apply comprehensive evaluation metrics, including four regression metrics - Spearman correlation coefficient (SCC), Pearson’s correlation coefficient (PCC), root mean square error (RMSE) and mean absolute error (MAE) - as well as three multi-class classification metrics: accuracy (ACC), Matthews correlation coefficient (MCC) and Cohen’s kappa score (*κ*). Higher SCC, PCC, ACC, MCC and *κ* values indicate superior performance, whereas lower RMSE and MAE values correspond to smaller prediction errors (see “Methods” for more details).

Table 1 presents the performance of various methods on the S461 dataset. DeepTherm achieved the lowest errors, with an RMSE of 0.926 and an MAE of 0.671, compared with the best values among the other methods (RMSE 0.984; MAE 0.722; Table S1 in the ESM). It also achieved the highest SCC of 0.86 and PCC of 0.84. Figure 2a compares the absolute errors for individual mutations; the pairwise P-values are shown above the violins.

**Table 1.**
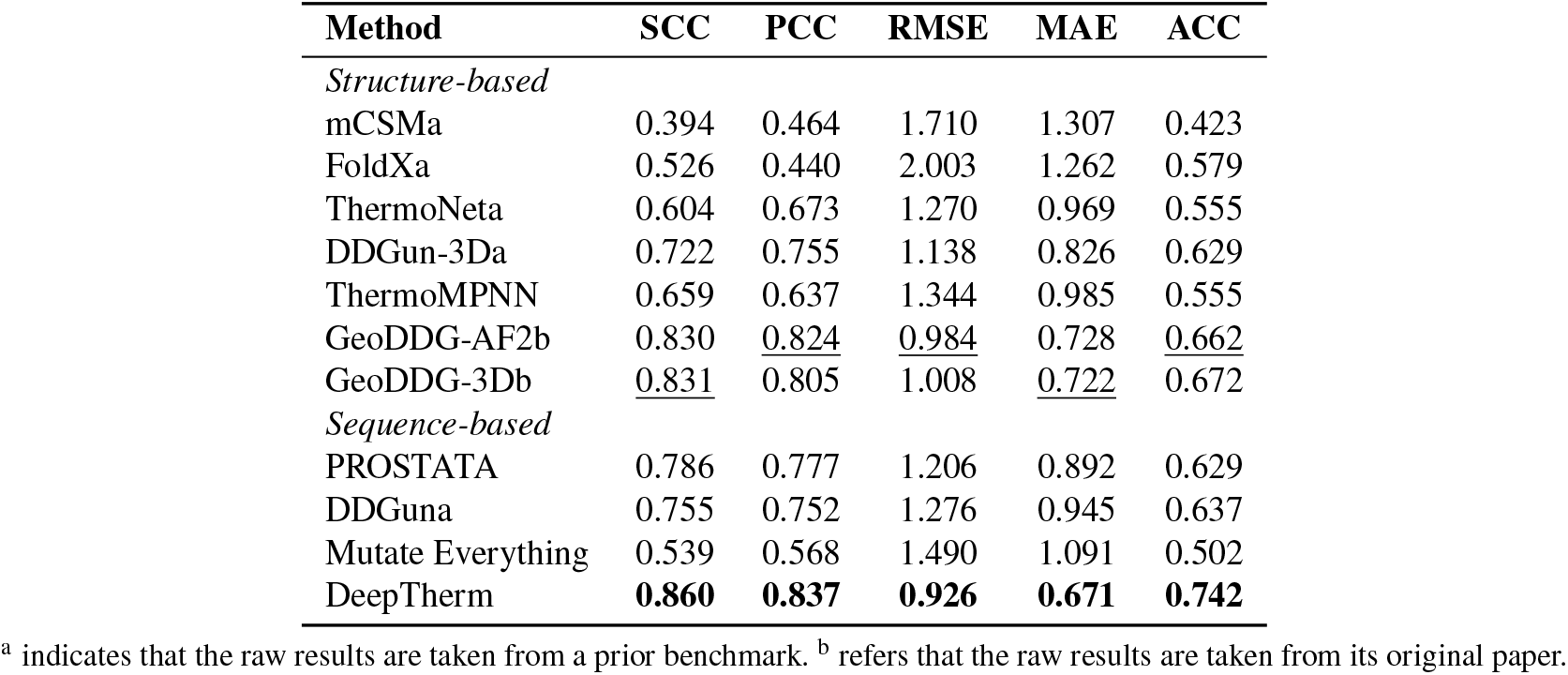
Overall comparison of ΔΔ*G* prediction on S461 in terms of SCC, PCC, RMSE, MAE, and ACC.

**Fig. 2.**
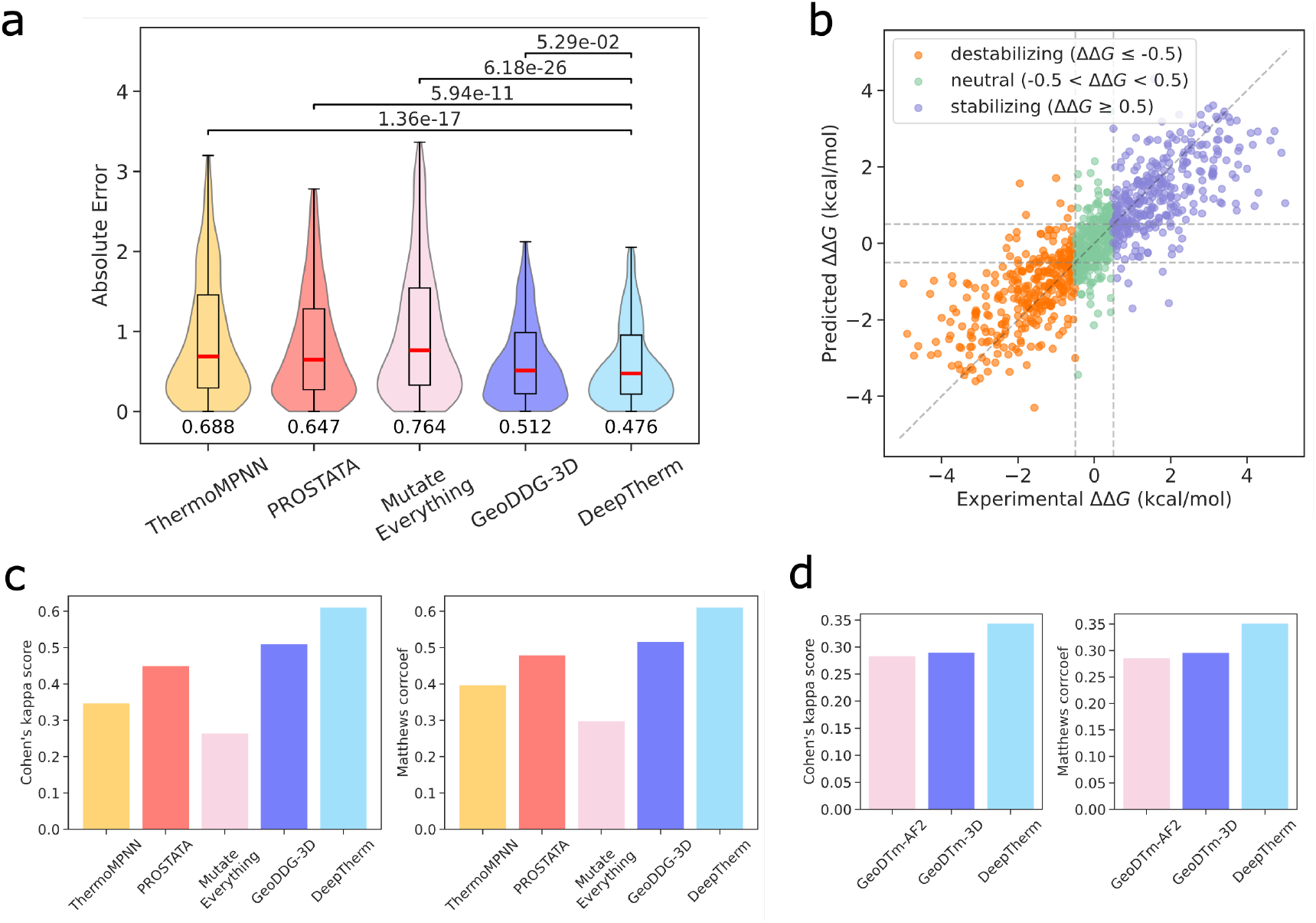
Performance comparison of DeepTherm. **a**. The violin plot of DeepTherm and other representative methods in terms of absolute error. The median values are denoted by the red lines. The first and third quartiles are indicated by the bounds of the box. The whiskers represent the 0.8 interquartile range (IQR). Two-sided paired t-tests are conducted between DeepTherm and other methods, and the P-values are annotated at the top of the violins. **b**. The detailed predictions of DeepTherm. The mutations are divided into three categories with different ΔΔ*G* thresholds. **c**. The performance of DeepTherm and other representative methods in terms of MCC and *κ* score for ΔΔ*G* prediction. **d**. The performance of DeepTherm and other representative methods in terms of MCC and *κ* score for Δ*T*_*m*_ prediction.

In certain scenarios such as soluble protein design, classifying mutations rather than predicting exact ΔΔ*G* values is more practically useful [33–35]. To establish a classification task, we categorize mutations into three classes: destabilizing (ΔΔ*G <* −0.5 kcal/mol), stabilizing (ΔΔ*G >* 0.5 kcal/mol) and neutral. DeepTherm achieves a significantly higher ACC than other methods (Table 1). Among the 922 mutations (comprising 461 direct and 461 inverse mutations), DeepTherm correctly predicts the mutation classes for 684 cases (Fig. 2b). Additionally, DeepTherm achieves the highest MCC and *κ* (both *>* 0.61), whereas the second-best method, GeoDDG-3D, achieves only 0.51 (Fig. 2c), highlighting its superior performance in mutation classification.

Furthermore, we compare DeepTherm with GeoDTm-AF2 [16] and GeoDTm-3D [16] on Δ*T*_*m*_ prediction. Interestingly, DeepTherm achieves a comparable SCC to GeoDTm-AF2 and GeoDTm-3D while significantly outperforming them in terms of PCC, RMSE and MAE (Table 2 and Table S2 in the ESM). For classification, we similarly categorized mutations into three groups: destabilizing (Δ*T*_*m*_ *<* −2 °C), stabilizing (Δ*T*_*m*_ *>* 2 °C), and neutral. DeepTherm achieved an ACC of 0.561, compared with GeoDTm-AF2 (0.522) and GeoDTm-3D (0.525) (Table 2). Moreover, the evaluation on Cohen’s kappa score and MCC confirms the better performance of DeepTherm relative to the two GeoDTm models (Fig. 2d). Overall, our data indicates that DeepTherm achieves a stronger or comparable performance for both ΔΔ*G* and Δ*T*_*m*_ prediction.

**Table 2.**
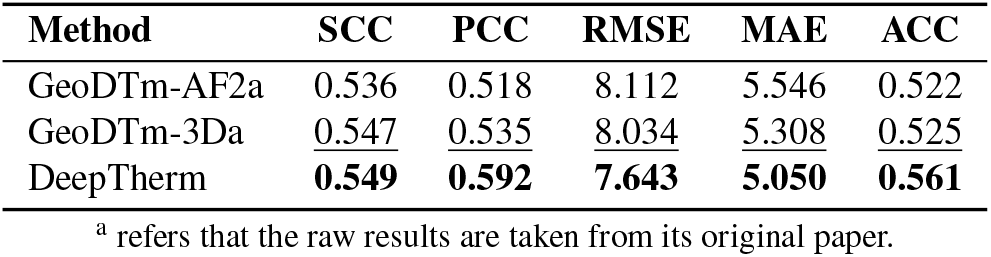
Overall comparison of Δ*T*_*m*_ prediction on S571 in terms of SCC, PCC, RMSE, MAE, and ACC.

### 3.4 DeepTherm provides accurate guidance on individual proteins

We evaluate DeepTherm’s performance on individual proteins to investigate its capability for protein engineering. We independently evaluate its performance on individual proteins in S461. Accordingly, there are 388 mutations from 15 proteins used here; each protein has at least 6 mutations. DeepTherm exhibits substantial improvements in SCC, PCC, and ACC, compared to GeoDDG-3D (Fig. 3a). Specifically, it achieves remarkable median improvements of 9.1% (0.88 versus 0.81) in SCC, 7.5% (0.86 versus 0.80) in PCC, and 19.4% (0.80 versus 0.67) in ACC. Regarding error ranges, DeepTherm surpasses GeoDDG-3D across most proteins (Fig. 3b). A similar conclusion can also be obtained for Δ*T*_*m*_ prediction (Fig. S1 in the ESM). Collectively, these results highlight the advantages of DeepTherm in ΔΔ*G* and Δ*T*_*m*_ at protein-level, which provides strong guidance for protein engineering.

**Fig. 3.**
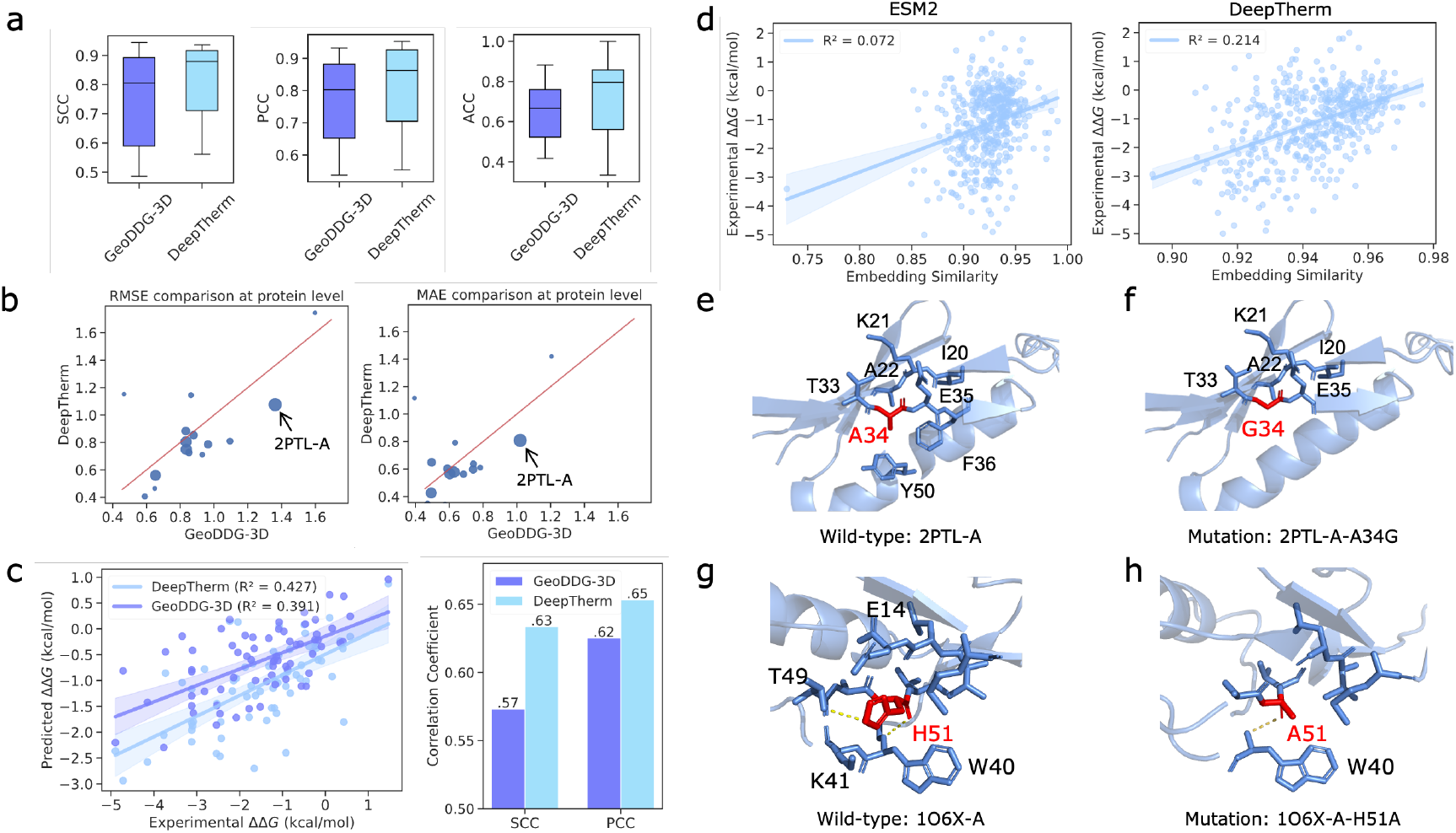
Performance comparison and analysis at protein-level. **a**. The performance comparison of DeepTherm and other methods on 15 individual proteins in terms of SCC, PCC and ACC. **b**. The performance comparison of DeepTherm and other methods on 15 individual proteins in terms of RMSE and MAE, where each point represents a protein, and the point size indicates the number of mutations. **c**. Comparison of detailed predictions of 2PTL-A by DeepTherm and other methods in terms of correlation. **d**. The correlation between representation similarity and experimental ΔΔ*G* in original ESM2 and DeepTherm, where the similarity is calculated by single mutations and wild-type protein 2PTL-A. **e**. Local structure of 2PTL-A near the mutation site Ala34, where the residues with annotations are contacted with the wild-type residue A34. **f**. Local structure of 2PTL-A mutation (A34G) near the mutation site Gly34, where the residues with annotations are contacted with the mutated residue G34. **g**. Local structure of 1O6X-A near the mutation site H51, where the yellow lines indicate the hydrogen bonds, and the residues with annotations are associated with changes before and after the mutation H51A. **h**. Local structure of 1O6X-A mutation (H51A) near the mutation site Ala51, where the yellow lines indicate the hydrogen bonds, and the residues with annotations are associated with changes before and after the mutation H51A.

We investigate protein 2PTL-A, which is a clear sample of DeepTherm outperforming GeoDDG-3D (Fig. 3b). Fig. 3c presents a head-to-head comparison of DeepTherm and GeoDDG-3D for ΔΔ*G* prediction on all known mutations of 2PTL-A, which indicate that DeepTherm achieves higher *R*^2^ correlation values with an improvement of 9% over GeoDDG-3D, as well as SCC (increase 11%) and PCC (increase 5%). These demonstrate that DeepTherm can provide a more reliable rank list for mutations of 2PTL-A. To explore the reason for the DeepTherm boost, we analyze the representation similarities of these mutations from our model and ESM2 that are also used in GeoDDG-3D. In the original ESM2 model, these mutations have high similarities and are hard to distinguish for ΔΔ*G* prediction, while in DeepTherm, the correlation between representation similarities and ΔΔ*G* is significantly improved (Fig. 3d).

DeepTherm is not only able to identify positive mutations but also has high accuracy to predict negative mutations, while current structure-based predictors often fail to reliably identify destabilizing mutations. For instance, GeoDDG-3D misclassifies destabilizing variants such as A34G in 2PTL-A and H51A in 1O6X-A as neutral or stabilizing. In contrast, DeepTherm accurately captures both cases, demonstrating its higher resolution in identifying negative mutations. As shown in Fig. 3e and 3f, A34G is a curated destabilizing mutation (ΔΔ*G* = −2.1 kcal/mol), which GeoDDG-3D predicts as nearly neutral (ΔΔ*G* = −0.3 kcal/mol), whereas DeepTherm outputs a substantially more accurate prediction (ΔΔ*G* = −1.2 kcal/mol). Structural analysis via FoldX reveals that replacing alanine with glycine at position 34 removes the *C*_*β*_ atom, reducing solvent accessible surface area (SASA) and local backbone flexibility. Furthermore, the smaller molecular volume of glycine weakens contacts with neighboring residues, particularly Y50 and the hydrophobic residue F36, thereby diminishing hydrophobic interactions and reducing local stability (Fig. 3f).

A similar pattern is observed for the H51A mutation in protein 1O6X-A (Fig. 3g-h). While DeepTherm accurately predicts its destabilizing effect (ΔΔ*G* = −0.8 kcal/mol), GeoDDG-3D incorrectly classifies it as stabilizing (ΔΔ*G* = 0.6 kcal/mol). In the wild-type structure, H51 forms hydrogen bonds with W40 and T49. The substitution with alanine significantly reduces side-chain volume, abolishes the hydrogen bond with T49, and disrupts packing interactions with E14 and K41. The loss of the aromatic ring further destabilizes the local structure by creating a cavity. These structural features, which underlie the decrease in stability (experimental ΔΔ*G* = −0.7 kcal/mol), are effectively captured by DeepTherm but overlooked by GeoDDG-3D.

### 3.5 Generalization of self-distillation inference and antisymmetric constraint in DeepTherm

The self-distillation inference strategy can enhance training data diversity and improve the accuracy of existing trained models. After removing the inference stage, the performance will reduce in all metrics (Fig. 4a). Specifically, for ΔΔ*G* prediction, both SCC and PCC reduce by 3%, and RMSE and MAE correspondingly increase by 6% and 4%. This phenomenon also exists in Δ*T*_*m*_ (Fig. 4b).

**Fig. 4.**
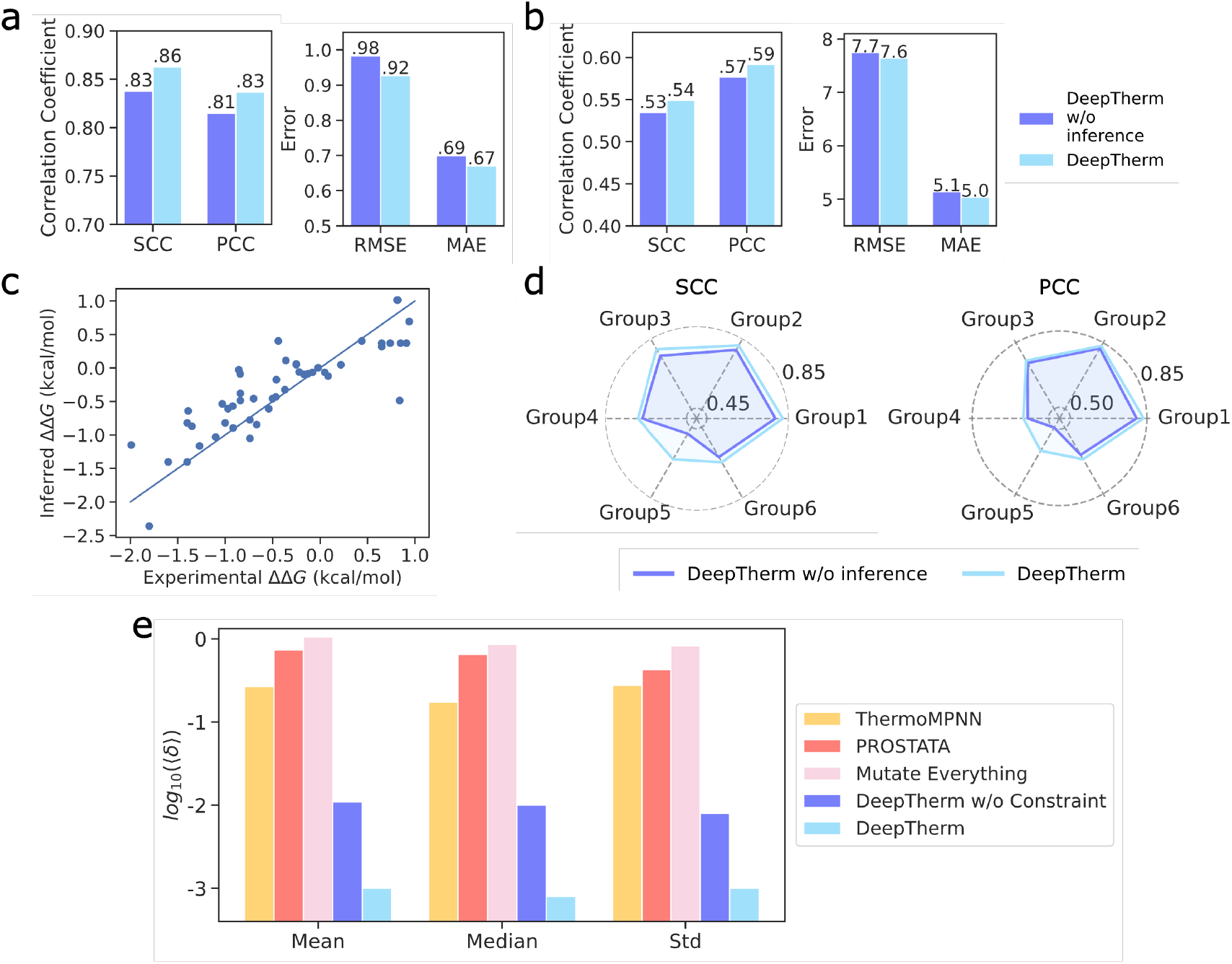
The role analysis of self-distillation strategies. **a**. The performance comparison of DeepTherm and DeepTherm without self-distillation strategies (inference stage), in correlation coefficient (left panel, including SCC and PCC), error ranges (right panel, RMSE and MAE) for ΔΔ*G* prediction. **b**. The performance comparison of DeepTherm and DeepTherm without self-distillation strategies (inference stage), in correlation coefficient (left panel, including SCC and PCC), error ranges (right panel, RMSE and MAE) for Δ*T*_*m*_ prediction. **c**. The correlation between experiment ΔΔ*G* and inferred ΔΔ*G* from cDNA display proteolysis. **d**. The performance comparison of DeepTherm and DeepTherm without self-distillation strategies (inference stage) on different test groups from cDNA display proteolysis data. The left panel and right panel indicate SCC and PCC, respectively. **e**. The antisymmetric error comparison for direct and inverse mutations.

To investigate the role of self-distillation more thoughtfully, we collect more protein stability-related data, which is generated by cDNA display proteolysis for fast measuring folding stability [30]. It infers Δ*G* of proteins and ΔΔ*G* of mutations from sequencing counts with a Bayesian model. To validate the accuracy of inferred ΔΔ*G* values, we extract the overlapping mutations in the inferred data and our training data. The correlation between them achieves 0.85 when considering ΔΔ*G* between −2 and 1 kcal/mol (Fig. 4c). Consequently, we split all mutations into six groups by protein length and randomly sample three proteins with 450 mutations in each group (Fig. S2 in the ESM), resulting in six sampled independent test datasets (Table S3 in the ESM). Then, based on our trained teacher models, we first generate pseudo-labels for these test datasets and re-train corresponding DeepTherm models, respectively. As shown in Fig. 4d, SCC and PCC values at the inference stage can be further improved in most cases, which demonstrates the effectiveness of self-distillation in our model.

Additionally, as can be deduced from the definitional formula of ΔΔ*G*_*wt→mut*_ = Δ*G*_*mut*_ − Δ*G*_*wt*_, direct and inverse mutations should produce exactly opposite ΔΔ*G* values. Since DeepTherm does not employ a fully symmetrical network architecture (i.e., GeoDDG-AF2 and GeoDDG-3D), it may not produce inverse outputs for symmetric inputs (*x*_*wild*_, *x*_*mut*_) and (*x*_*mut*_, *x*_*wild*_). This limitation also exists in other methods, such as PROSTATA, ThermoMPNN, and Mutate Everything, which need to be addressed to make models more robust in practical cases. To mitigate this limitation, we introduce a constraint loss to guide our model in learning this antisymmetric property. The absolute value of the sum of predicted results for direct and inverse mutations is used here as an error evaluation metric ⟨*δ*⟩. It can be observed that after introducing the antisymmetric constraint, DeepTherm exhibits a significant reduction in error and is nearly capable of predicting perfectly opposite ΔΔ*G* values for direct and inverse mutations, making it more valuable for practical applications (Fig. 4e). Above all, the self-distillation strategy can improve the robustness of our model, and the antisymmetric constraint makes the predictions more practical.

## 4 Discussion

Accurately predicting protein stability changes (ΔΔ*G* and Δ*T*_*m*_) of mutations is crucial for protein design and protein engineering. Sequence-based predictors are more easily applicable to large-scale datasets compared to structure-based approaches. However, existing sequence-based methods fail to capture structural contact changes, resulting in inferior performance to structure-based methods. In this study, we develop a sequence-based deep learning method for predicting protein stability changes from single residue mutations, called DeepTherm. It leverages a pLLM to encode residue representations and predict residue contacts. Subsequently, a cross-attention mechanism is introduced to capture the contact changes around the mutated positions. Comprehensive and rigorous analyses highlight the significant improvements achieved by DeepTherm. Notably, DeepTherm not only demonstrates strong overall performance, but also shows its advantages at the individual protein level, which is more efficient in practical cases.

Although fine-tuning pLLMs for predicting protein stability changes has already been explored in previous studies [22, 23], these models still face the limitations of ΔΔ*G* training data, which is very unbalanced. This can lead to decreased performance of these models when predicting the independent test set with different data distributions [35]. Consequently, we adopt a self-distillation strategy at the inference stage to improve the generalization of models. This strategy can generate more data and enrich data distributions, which further improves the performance of existing models. On the other hand, traditional asymmetrical frameworks cannot generate inverse outputs for symmetric inputs, except when the wild-type and mutated proteins share the whole network. However, completely sharing model weights between wild-type and mutated proteins may make it difficult to learn the difference between these two proteins. To solve these limitations, we introduce a simple antisymmetric constraint to guide our model to learn antisymmetric data. Comprehensive ablation studies suggest that the self-distillation strategy substantially enhances the performance of DeepTherm, offering new perspectives into improving existing protein large language models (pLLMs) for specific downstream tasks. Contrastive loss can also enable an asymmetric network architecture to better learn the constraints of antisymmetric data.

## 5 Conclusions

Overall, DeepTherm only takes protein sequences as input and does not rely on high-quality structural data. It can predict residue contact changes and potentially capture the key residues or regions involved in important interactions that maintain protein stability, such as *π* − *π* interactions and hydrogen bonds. In the future, we will consider side-chain changes and introduce more geometric potential to help models capture structural changes during mutations. Additionally, in practical protein engineering, optimizing a protein for specific applications often requires multiple mutations. Consequently, how to design an effective framework to predict stability changes upon multiple residue mutations is another challenge to be addressed.

Electronic Supplementary Material: Supplementary Figures S1–S2 and Tables S1–S3 are provided in the accompanying ESM file.

## Acknowledgements

This work is supported in part by the National Key Research and Development Program of China (2023YFF1205500 to R.Z.), the National Natural Science Foundation of China (No. 625B2184 to W.W. and No. 62225209 to M.L.), and the High Performance Computing Center of Central South University.

## Availability of data and materials

The data and source code supporting this study are available at https://github.com/CSUBioGroup/DPStab.

## Author biographies

**Wenkang Wang** Wenkang Wang received the B.S. degree from Central South University, China, in 2020. He is working toward the Ph.D. degree in computer science and engineering at Central South University, China. His research interests include bioinformatics, complex network analysis, machine learning and deep learning.

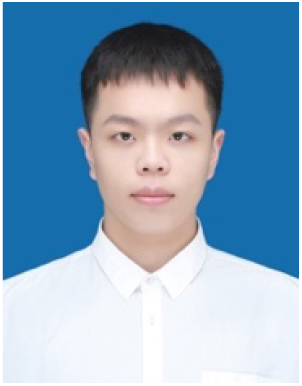

**Min Li** Min Li received the Ph.D. degree in computer science from Central South University, China, in 2008. Currently, she is a professor and dean of the School of Computer Science and Engineering, Central South University, China. Her main research interests include bioinformatics and systems biology. She has published more than 100 technical papers in refereed journals such as Cell, Nature Communications, Genome Research, Bioinformatics, Briefings in Bioinformatics, IEEE/ACM Transactions on Computational Biology and Bioinformatics, and conference proceedings such as BIBM, GIW and ISBRA. According to Google Scholar, her publications have received more than 20,000 citations, and her h-index is 70.

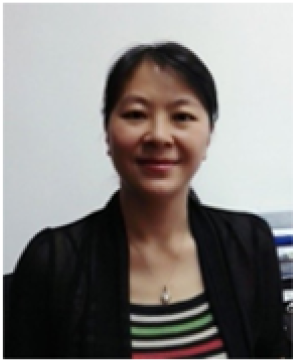

