## Supplementary Files for "Self-distillation and antisymmetric constraints enable accurate prediction of protein stability changes from single mutations"

### Electronic Supplementary Material

**This PDF file includes:**

Figures S1 to S2
Tables S1 to S3

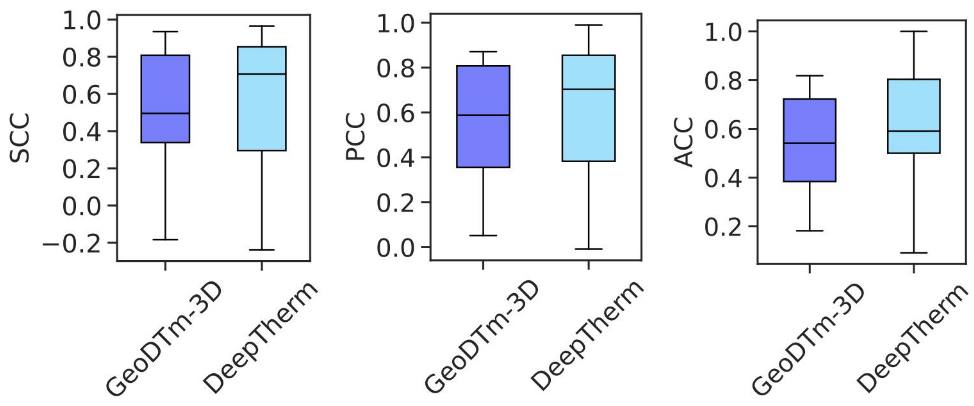


Fig. S1. The performance comparison of DeepTherm and GeoDTm-3D on 15 individual proteins in terms of SCC, PCC and ACC.

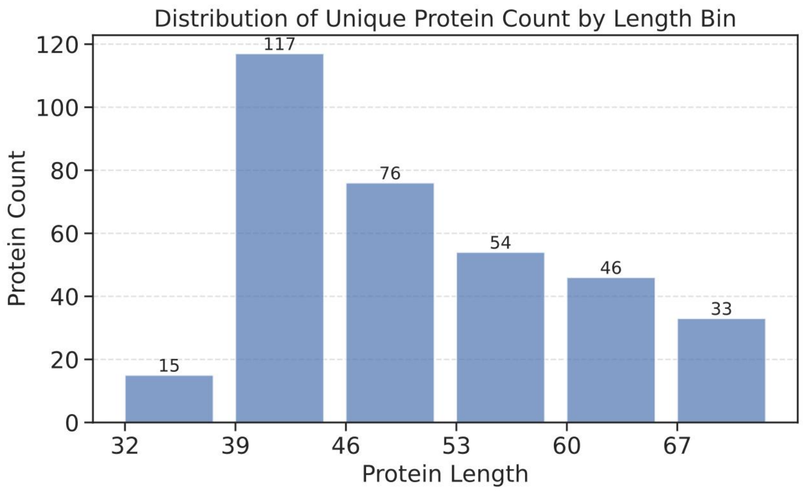

Fig. S2. The distribution of proteins generated by cDNA display proteolysis data.

Table S1. Detailed comparison of $\boldsymbol{\Delta\Delta G}$ prediction on S461 in terms of SCC, PCC, RMSE, MAE, and ACC.

| **Method** | **Direct** | | | | | **Inverse** | | | | |
|---|---|---|---|---|---|---|---|---|---|---|
| | **SCC** | **PCC** | **RMSE** | **MAE** | **ACC** | **SCC** | **PCC** | **RMSE** | **MAE** | **ACC** |
| mCSM[a] | 0.514 | 0.535 | 1.070 | 0.803 | 0.67 | 0.123 | 0.176 | 2.168 | 1.812 | 0.171 |
| FoldX[a] | 0.349 | 0.224 | 2.232 | 1.382 | 0.601 | 0.447 | 0.407 | 1.743 | 1.141 | 0.557 |
| ThermoNet[a] | 0.487 | 0.556 | 1.228 | 0.925 | 0.555 | 0.430 | 0.516 | 1.311 | 1.013 | 0.555 |
| DDGun-3D[a] | 0.583 | 0.636 | 1.101 | 0.800 | 0.629 | 0.551 | 0.587 | 1.174 | 0.851 | 0.629 |
| ThermoMPNN | 0.453 | 0.456 | 1.280 | 0.927 | 0.607 | 0.466 | 0.480 | 1.406 | 1.044 | 0.503 |
| GeoDDG-AF2[b] | <u>0.682</u> | 0.690 | 0.983 | 0.728 | 0.662 | <u>0.682</u> | 0.690 | <u>0.984</u> | 0.728 | <u>0.662</u> |
| GeoDDG-3D[b] | 0.669 | 0.668 | 1.008 | 0.722 | 0.672 | 0.669 | 0.668 | 1.008 | <u>0.722</u> | 0.672 |
| PROSTATA | 0.656 | 0.664 | <u>0.961</u> | <u>0.697</u> | <u>0.716</u> | 0.533 | 0.566 | 1.410 | 1.086 | 0.542 |
| DDGun[a] | 0.569 | 0.588 | 1.251 | 0.939 | 0.644 | 0.546 | 0.551 | 1.302 | 0.951 | 0.629 |
| Mutate Everything | 0.630 | 0.622 | 1.035 | 0.752 | 0.681 | 0.135 | 0.164 | 1.835 | 1.429 | 0.323 |
| DeepTherm | 0.690 | <u>0.680</u> | 0.926 | 0.671 | 0.742 | 0.690 | <u>0.679</u> | 0.926 | 0.671 | 0.742 |

[a] indicates that the raw results are taken from a prior benchmark. [b] refers that the raw results are taken from its original paper.

Table S2. Detailed comparison of $\boldsymbol{\Delta T_m}$ prediction on S571 in terms of SCC, PCC, RMSE, MAE, and ACC.

| **Method** | **Direct** | | | | | **Inverse** | | | | |
|---|---|---|---|---|---|---|---|---|---|---|
| | **SCC** | **PCC** | **RMSE** | **MAE** | **ACC** | **SCC** | **PCC** | **RMSE** | **MAE** | **ACC** |
| GeoDTm-AF2[a] | 0.513 | 0.462 | 8.112 | 5.546 | 0.522 | 0.513 | 0.462 | 8.112 | 5.546 | 0.522 |
| GeoDTm-3D[a] | 0.510 | <u>0.467</u> | <u>8.034</u> | <u>5.308</u> | <u>0.525</u> | 0.510 | <u>0.467</u> | <u>8.034</u> | <u>5.308</u> | <u>0.525</u> |
| DeepTherm | <u>0.512</u> | 0.526 | 7.642 | 5.050 | 0.562 | <u>0.512</u> | 0.526 | 7.643 | 5.050 | 0.560 |

[a] refers that the raw results are taken from its original paper.

Table S3. The details of proteins sampled from cDNA display proteolysis data.

| Group | Protein | Length | Mutations |
|---|---|---|---|
| 1 | 2M9F | 33 | 150 |
| 1 | 2M9I | 34 | 150 |
| 1 | 1WR4 | 36 | 150 |
| 2 | HHH_rd1_0473 | 43 | 150 |
| 2 | r4_412_TrROS_Hall | 45 | 150 |
| 2 | HEEH_KT_rd6_1415 | 43 | 150 |
| 3 | 4G3O | 47 | 150 |
| 3 | EA\|run2_0325_0005 | 47 | 150 |
| 3 | EA\|run6_0680_0006 | 47 | 150 |
| 4 | 2EXD | 58 | 150 |
| 4 | 2LP5 | 57 | 150 |
| 4 | 1Y0M | 58 | 150 |
| 5 | 2OCH | 62 | 150 |
| 5 | 6M3N | 64 | 150 |
| 5 | 2MCH | 64 | 150 |
| 6 | 1ZHC | 68 | 150 |
| 6 | 2KCM | 68 | 150 |
| 6 | 7BPM | 69 | 150 |
